# Antifungal activity of Propranolol against *Fusarium* keratitis isolates from the Mycotic Ulcer Treatment Trial (MUTT) and the United States

**DOI:** 10.1101/2021.01.19.427292

**Authors:** Ruina Bao, Brandon S. Ross, Cecilia G. Perez, Galini Poimenidou, N. Venkatesh Prajna, Robert A. Cramer, Michael E. Zegans

**Author notes:** Address correspondence to Michael E. Zegans. M.E.Z. and R.A.C. contributed equally to this study. RB and BSR contributed equally to this study.

## Abstract

**Background:** *Fusarium* keratitis is an infection of the cornea that often results in corneal perforation requiring corneal transplantation even with topical ocular antifungal therapy. The polyene natamycin remains the current antifungal of choice for *Fusarium* keratitis, but prompt sterilization of the cornea is often not achieved with contemporary therapy. Recently, natamycin synergy with the beta-adrenergic antagonist timolol against *Fusarium* species was reported.

**Objective:** Our objective in this study was to characterize the *in vitro* antifungal effects of additional beta-adrenergic antagonists alone or in combination with natamycin on *Fusarium* keratitis isolates from the Mycotic Ulcer Treatment Trial (MUTT) and USA.

**Methods:** Microbroth dilution assays were used to determine the minimal inhibitory concentration (MIC) of beta-adrenergic antagonists against 18 *Fusarium spp*. keratitis (10 from MUTT, 8 from USA) and 3 *Aspergillus fumigatus* isolates. The fractional inhibitory concentration index (FICI) was calculated to assess interactions with natamycin.

**Results:** Most beta-blockers did not show antifungal activity or synergy with natamycin with the exception of propranolol. A racemic mix of propranolol had fungicidal activity with MIC between 31 and 83 μg/mL for the *Fusarium* isolates. The MIC of the less cardioactive R enantiomer was lower (27-83 μg/mL) than the MIC of the S enantiomer (42-104 μg/mL). The MICs of both propranolol and natamycin were lower in combination but were not synergistic. The MIC of propranolol was 156 μg/mL for the *A. fumigatus* isolates.

**Conclusions:** Propranolol has intrinsic *in vitro* fungicidal activity and lowers the MIC of natamycin. Both the R and S enantiomers of propranolol had antifungal activity with the MIC modestly but significantly lower for R-propranolol. These findings have relevance both for the treatment of fungal keratitis and of glaucoma in the setting of fungal keratitis. Further study of propranolol’s antifungal activity may lead to a novel treatment for fungal keratitis and possibly other fungal infections.

**Trial Registration:** ClinicalTrials.gov Identifier: NCT00997035 (MUTT Trial)

## INTRODUCTION

Fungal keratitis (FK), most commonly caused by the genera *Fusarium*, is a severe and often blinding infection.^1,2^ Currently, there is only one Food and Drug Agency (FDA) approved antifungal for FK, the polyene natamycin.^3^ The Mycotic Ulcer Treatment Trials (MUTT-I and II) explored the role of topical and/or oral voriconazole in the treatment of fungal keratitis.^4,5^ MUTT-I was halted due to excess perforations and corneal transplantation requirements in the voriconazole-treated group. MUTT-II was unable to show a statistically significant value of adjunctive oral voriconazole compared to placebo in a group of more severe ulcers. Thus, voriconazole did not have advantages over natamycin and there remains a need for novel antifungal strategies to treat fungal keratitis. Equally important, both studies found a high rate of persistently positive cultures at 6 days leading to high rates of corneal perforation and transplantation (16% in MUTT-I and 51% in MUTT-II).^4–7^ Cultures of the excised corneal buttons from transplantation cases in MUTT-II were positive for fungi in 67% (45/67) of cases.^8^ Taken together these data indicate that there is often a failure to obtain a microbiologic cure of the cornea even with natamycin and oral voriconazole treatments, and this persistence of cultivable fungi in the face of treatment is associated with poor clinical outcomes.

We recently reported that the beta-adrenergic antagonist (a.k.a. beta blocker) timolol has synergistic antifungal activity with natamycin against filamentous fungi at concentrations comparable to those used for the treatment of glaucoma.^9^ However, timolol did not have intrinsic antifungal activity as a single agent. In this study we describe our discovery that propranolol has intrinsic fungicidal activity against *Fusarium* species keratitis isolates at concentrations achievable in the cornea.

## MATERIALS AND METHODS

### Strains and Culture Conditions

*Fusarium solani* strains 06-0110, 06-0111,06-0133, 06-330, and 06-0487 were obtained from the University of California at San Francisco-Proctor Foundation, DUMC 132.02 is from Duke University Medical Center (gift of Dr. Wiley Schell). *Fusarium oxysporum* strains 06-0197 and 06-0342 were also gifts from UCSF. Ten *Fusarium solani* keratitis isolates from the Mycotic Ulcer Treatment Trial (MUTT) were randomly selected using random.org. Frozen glycerol stocks of microconidia were streaked onto potato dextrose agar (PDA; Becton, Dickinson, Franklin Lakes, NJ) and incubated for 72 hours at 30°C in atmospheric conditions. An agar plug containing hyphae was used to inoculate 100mL of potato dextrose broth (PDB; Becton, Dickinson) in a baffled culture flask. Shaking cultures were grown at 30 °C for 72 hours at 200 rpm. Microconidia were harvested by filtering liquid cultures through a sterile funnel lined with Miracloth. Filtrates were centrifuged at 5000 rpm for 5 min at 4°C, decanted and washed with phosphate buffered saline (PBS; Corning, Mediatech Inc., Manassas, VA). Conidia were counted using a hemocytometer and the inoculum was resuspended in RPMI 1640 (Gibco RPMI 1640 Medium #11875176 ThermoFisher Scientific, Grand Island, NY) at 1×10^5^ conidia/mL.

### *In vitro* antifungal activity of beta-blockers

Compounds R-, R/S- and S-propranolol hydrochloride (Sigma-Aldrich, St. Louis, MO) or nadolol (Sigma-Aldrich), R/S-atenolol, and sotalol hydrochloride (Tocris Bioscience, Minneapolis, MN) were dissolved in RPMI 1640 to a stock concentration of 1 mg/mL or 5mg/ml, respectively. 100 μL of the stock was added to the first well on a 96 well plate and serial 2-fold dilutions with RPMI were used to fill each row. 100 μL of conidial suspension were added to each well and plates were incubated at 30 °C for 72 hours at atmospheric conditions, or at 37°C for 24-48 hours at 5% CO_2_ for *A. fumigatus* strains. The minimum inhibitory concentration (MIC) was determined by viewing individual wells under the light microscope and set for the drug concentration which had minimal to no germination of the microconidia.

Stock solutions of natamycin (10 mg/mL, Sigma-Aldrich, St. Louis, MO) were solubilized in DMSO. Serial dilutions were prepared by dissolving this stock solution into RPMI-1640 media. In the checkerboard assay, 50 μL of RPMI + natamycin was added to each well of row A in a 96 well plate. 50 μL of RPMI + propranolol was added to each well of column A. 50 μL of decreasing two-fold dilutions were added either from top to bottom (natamycin) or left to right (propranolol). The last solution of each dilution series was 0 μg/mL. *Fusarium* microconidia were diluted to a concentration of 1×10^5^ conidia/mL in 10 mL of RPMI-1640, and 100 μL of the inoculum was added to each well. The fractional inhibitory concentration index (FICI) of the combination was determined for each isolate according to previously published methods.^9^

## RESULTS

Propranolol was observed to have antifungal activity against *Fusarium* and *Aspergillus* isolates (Table 1). At the MIC, all three formulations inhibited *Fusarium* microconidia germination *in vitro* as observed under DIC microscopy (Figure 1A-B). Similar findings were observed in *A. fumigatus* isolates, but the MICs were two-fold higher at 156 μg/ml (Table 1). When microconidia treated with MIC concentrations of drug were plated onto PDA, no fungal growth was detected after incubation at 30 °C for 72 hours for all 18 *Fusarium* isolates, in contrast to the robust growth observed in untreated conidia (Figure 1C-D). These data suggest propranolol has fungicidal activity against *Fusarium* microconidia. Natamycin decreased the MIC of propranolol in 16 of 18 (89%) of isolates with 11 of 18 showing more than a twofold reduction (Table S1). The fractional inhibitory concentration indices (FICI) were all ≥0.5 denoting an indifferent interaction between natamycin and propranolol, though the MIC of both compounds was reduced in combination.^10^

**Figure 1:**
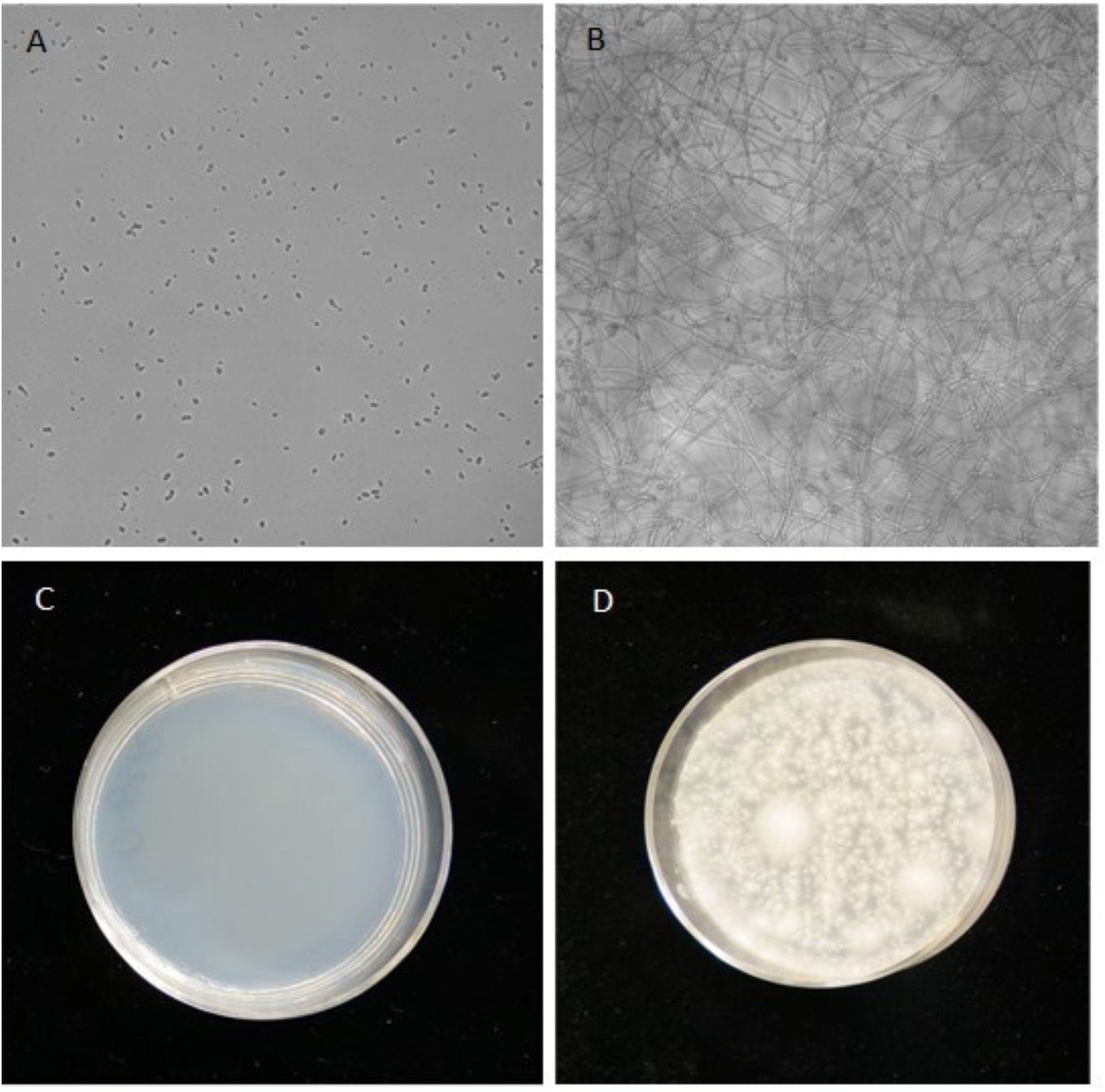
Representative images of fungal growth viewed under differential interference contrast microscopy. A) *Fusarium solani* 06-0110 microconidial germination was inhibited with 63 μg/mL of propranolol dissolved in RPMI with L-glutamine and sodium bicarbonate after incubation at 30 °C for 72 hours in atmospheric conditions. B) DIC image of the untreated positive control shows abundant mycelial growth. C) Representative images of keratitis isolates from American and Indian patients show growth after contents of wells from plates used for the CLSI microbroth dilution method of determining minimum inhibitory concentrations. 06-0110 *Fusarium solani* is shown here. Wells containing the MIC concentrations of propranolol (31-63 μg/mL) exhibited no growth when plated. D) The untreated, positive control wells exhibited robust growth after incubating for 72 hours at 30 °C in atmospheric conditions. Three biological replicates were performed for each isolate.

**Table 1:** Propranolol has antifungal activity against *Fusarium* keratitis isolates as well as *Aspergillus* isolates.

| Isolate | Natamycin<br>MIC<sup>a,b</sup><br>(µg/mL) | R-propranolol<br>MIC<sup>a,b</sup><br>(µg/mL) | S-propranolol<br>MIC<sup>a,b</sup><br>(µg/mL) | R/S-propranolol<br>MIC<sup>a,b</sup><br>(µg/mL) | Sotalol<br>MIC<sup>a,c</sup><br>(µg/mL) | Nadolol<br>MIC<sup>a,c</sup><br>(µg/mL) | Atenolol<br>MIC<sup>a,c</sup><br>(µg/mL) |
| --- | --- | --- | --- | --- | --- | --- | --- |
| MUTT1090Z | 8 | 83 | 104 | 83 | - | - | - |
| MUTT1121U | 16 | 52 | 52 | 31 | - | - | - |
| MUTT1146N | 32 | 52 | 52 | 63 | - | - | - |
| MUTT1170W | 21 | 27 | 42 | 42 | - | - | - |
| MUTT1584Q | 32 | 52 | 63 | 52 | - | - | - |
| MUTT1587W | 32 | 47 | 42 | 42 | - | - | - |
| MUTT1736Z | 16 | 31 | 42 | 42 | - | - | - |
| MUTT1753Z | 16 | 31 | 42 | 31 | - | - | - |
| MUTT2725Z | 16 | 42 | 63 | 52 | - | - | - |
| MUTT2226R | 27 | 47 | 52 | 52 | - | - | - |
| USA06-0110 | 27 | 42 | 52 | 52 | - | - | - |
| USA06-0111 | 21 | 26 | 52 | 42 | >2500 | >2500 | >2500 |
| USA06-0133 | 27 | 26 | 52 | 31 | >2500 | >2500 | >2500 |
| USA06-0197 | 16 | 42 | 52 | 52 | - | - | - |
| USA06-0342 | 27 | 42 | 63 | 52 | - | - | - |
| USA06-0330 | 27 | 26 | 52 | 42 | >2500 | >2500 | >2500 |
| USA06-0487 | 21 | 52 | 63 | 31 | >2500 | >2500 | >2500 |
| USA132.02 | 11 | 31 | 73 | 73 |  |  |  |
| Mean (SD) | 22 (7) | 43 (10) | 54 (9) | 48 (11) | >2500<br>(0) | >2500<br>(0) | >2500<br>(0) |
| - |  |  |  |  |  |  |  |
| CEA10 (*A. fumigatus*) | 48 | - | - | 156 | >2500 | >2500 | >2500 |
| AF293 (*A. fumigatus*) | 48 | - | - | 156 | >2500 | >2500 | >2500 |
| IFISW-F4 (*A. fumigatus*) | 48 | - | - | 156 | >2500 | >2500 | >2500 |
| Mean (SD) | 48 (0) | - | - | 156 (0) | >2500<br>(0) | >2500<br>(0) | >2500<br>(0) |

Abbreviations: MIC = minimum inhibitory concentration
<sup>a</sup> MICs rounded to whole numbers
<sup>b</sup> Average of data from three biological replicates
<sup>c</sup> Average of data from two biological replicates

## DISCUSSION

Our data suggest that propranolol has *in vitro* fungicidal activity against *Fusarium* species microconidia across a spectrum of corneal isolates from the MUTT study and from the USA. Similar to natamycin, most *Fusarium spp*. isolates in this study had an R propranolol MIC which was ≤50 μg/mL (Table 1). Furthermore, both the MIC of natamycin and propranolol were reduced when used in combination (Table S1). Though it did not achieve an FICI <0.5, the criteria for a synergistic drug interaction, a reduction in the MIC of natamycin is likely still of clinical relevance given that natamycin alone often fails to clear fungi from the cornea. Importantly, both enantiomers of propranolol had similar effects, suggesting use of the less cardio-active R enantiomer might allow increased dosing and therefore achieve higher drug levels in the cornea. The fact that both enantiomers are active as well as our inability to find *in silico* evidence of fungal beta-adrenergic receptors suggests that propranolol’s antifungal effects are not linked to the three canonical beta-receptors.^11^ Lastly, topical beta-blockers, including propranolol, have been safely used on the ocular surface at concentrations predicted to achieve an antifungal effect in the cornea furthers the clinical significance of this finding.^12^ Additional work to elucidate the mechanism of action of propranolol’s antifungal activity may reveal related compounds with even more potent antifungal activity or novel antifungal drug targets. This may prove to also have relevance for systemic infection with other filamentous fungi.

## TRANSPARENCY DECLARATIONS

None to declare

---

**Table S1.** R-propranolol and natamycin have additive to indifferent interactive effects against *Fusarium* keratitis isolates.

| Isolate | Natamycin MIC<sup>a</sup> (µg/mL) | R-Propranolol MIC<sup>a</sup> (µg/mL) | Natamycin Combination MIC<sup>a</sup> (µg/mL) | R-Propranolol Combination MIC<sup>a</sup> (µg/mL) | FICI<sup>b</sup> |
| --- | --- | --- | --- | --- | --- |
| MUTT1090Z | 8 | 125 | 4 | 63 | 1 |
| MUTT1121U | 16 | 31 | 8 | 4 | 0.63 |
| MUTT1146N | 16 | 63 | 8 | 8 | 0.63 |
| MUTT1170W | 32 | 31 | 16 | 8 | 0.75 |
| MUTT1584Q | 32 | 31 | 8 | 8 | 0.50 |
| MUTT1587W | 16 | 31 | 8 | 4 | 0.63 |
| MUTT1736Z | 8 | 31 | 8 | 4 | 0.63 |
| MUTT1753Z | 16 | 31 | 8 | 4 | 0.63 |
| MUTT2725Z | 16 | 31 | 8 | 8 | 0.75 |
| MUTT2226R | 16 | 31 | 8 | 4 | 0.63 |
| USA06-0110 | 32 | 31 | 4 | 16 | 0.63 |
| USA06-0111 | 16 | 16 | 8 | 8 | 1 |
| USA06-0133 | 16 | 16 | 8 | 8 | 1 |
| USA06-0197 | 16 | 31 | 8 | 16 | 1 |
| USA06-0330 | 16 | 16 | 8 | 8 | 1 |
| USA06-0342 | 32 | 31 | 4 | 16 | 0.63 |
| USA06-0487 | 16 | 31 | 4 | 16 | 0.75 |
| USA132.02 | 8 | 31 | 8 | 31 | 2 |

Abbreviations: MIC = minimum inhibitory concentration, FICI = fractional inhibitory concentration index
<sup>a</sup> MICs rounded to whole numbers.
<sup>b</sup> Values represent the average of data from 3 biological replicates, performed in one representative assay (<0.5 indicates synergistic interaction).

